# Post-encoding reactivation promotes one-shot learning of episodes in humans

**DOI:** 10.1101/2021.04.13.439658

**Authors:** Xiongbo Wu, Xavier Viñals, Aya Ben-Yakov, Bernhard P. Staresina, Lluís Fuentemilla

**Affiliations:** Cognition and Brain Plasticity Group, Bellvitge Institute for Biomedical Research, Hospitalet de Llobregat 08907, Spain; Department of Cognition, Development and Educational Psychology, University of Barcelona, Barcelona 08035, Spain; Institute of Neurosciences, University of Barcelona, Barcelona 08035, Spain; High School of Health Science, Tecnocampus Pompeu Fabra University, 08302 Mataró (Barcelona), Spain; Medical Research Council Cognition and Brain Sciences Unit, University of Cambridge, Cambridge CB2 7EF, United Kingdom; Department of Experimental Psychology, University of Oxford, UK

**Keywords:** Episodic memory, EEG, Memory reactivation, Pattern similarity, Encoding

## Abstract

Prior animal and human work have shown that post-encoding reinstatement plays an important role in organizing the temporal sequence of unfolding episodes in memory. Here, we investigated whether post-encoding reinstatement serves to promote the encoding of ‘one-shot’ episodic learning beyond the temporal structure in humans. In experiment 1, participants encoded sequences of pictures depicting unique and meaningful episodic-like events. We used representational similarity analysis on scalp electroencephalography recordings during encoding and found evidence of rapid picture elicited EEG patterns reinstatement at episodic offset (around 500ms post-episode). Memory reinstatement was not observed between successive elements within an episode and the degree of memory reinstatement at episodic offset predicted later recall for that episode. In experiment 2, participants encoded a shuffled version of the picture sequences from experiment 1, rendering each episode meaningless to the participant but temporally structured as in experiment 1, and we found no evidence of memory reinstatement at episodic offset. These results suggest that post-encoding memory reinstatement is akin to the rapid formation of unique and meaningful episodes that unfold over time.

## Introduction

In episodic encoding, an experienced event is rapidly transformed into a memory trace that has the potential to be consciously recollected at long-term (Tulving, 1983). Prior research has largely focused on examining how the brain contributes to successful encoding of individual trial information, such as single images (Paller and Wagner, 2002) or single item-context associations (Davachi, 2006). However, in natural settings, an episode is better characterized by a collection of successive elements that become contextually meaningful as they unfold over time. To be accessible for future retrieval, these elements have to be associatively linked into a bound memory trace. Discerning “when” the brain binds the continuous experience input into a cohesive episodic memory trace and “how” the brain undergoes this rapid transformation is essential to understand memory formation.

Human neuroimaging studies examining memory formation during a continuous stream of stimuli, such as naturalistic video clips, have shown that a distributed network of brain regions comprising the hippocampus and neocortex increased activity at the end of an event (Ben-Yakov et al., 2013; Ben-Yakov and Henson, 2018; Baldassano et al., 2017). This event offset brain signal has been shown to reflect a binding operation of the just encoded event elements into a specific spatio-temporal context (Ritchey and Cooper, 2020), which aligns well with the notion that the hippocampus is crucial for binding elements of our experience with contextual information (Davachi, 2006; Diana et al., 2007; Eichenbaum et al., 2007; Ranganath, 2010). In real life, episodic encoding relies on the possibility to form a coherent memory trace that integrates the temporally evolving sequence of elements into a meaningful context, so that if there is a shift in contextual information this is perceived as the end of one episode and the beginning of another (Zacks et al., 2007). These episodic boundaries are thought to support the segmentation of the continuous experience into discrete episodes (Zacks et al., 2007) and their detection has a direct impact on how events are organized into meaningful units in long-term memory (Kurby and Zacks, 2008; Radvansky, 2012; Ezzyat and Davachi, 2011; DuBrow and Davachi, 2013, 2014).

Work in rodents has provided evidence that memory replay at event offset plays a critical role in stabilizing a temporal memory organization beyond initial learning processes (Foster and Wilson, 2006; Diba and Buzsáki, 2007; Karlsson and Frank, 2008; Carr et al., 2011). In humans, rapid event offset memory reinstatement has been shown to be induced at the detection of context shifts during encoding of sequences of episodes and to predict their temporal order memory accuracy of the encoded sequential episodes in a later memory test (Sols et al., 2018; Silva et al., 2019). These findings suggest that the reactivation of an event contemporaneously with the experience of a subsequent adjacent event could theoretically result in the co-activation of the past and present events, promoting the binding of sequential events in their temporal order. In the real world, however, the recall of encoded episodes does not always depend on maintaining the exact order of the sequential representations, which can be fragile in many situations. Instead, it has been shown that when individuals are asked to recall episodes encoded in naturalistic conditions, they structured the recall along the causal (Brownstein & Read, 2007), semantics (van Kesteren et al., 2013; Baldassano et al., 2018) and the relations between the elements embedded in the episodes (Lee and Chen, 2021). In fact, psychological models of event comprehension have emphasized that as the experience unfolds, memory is carved by how people construct a coherent model of a situation, which consists of agents and objects, semantic and spatiotemporal contexts, and the relations between them (Radvansky and Zacks, 2011).

Notably, neuroimaging studies using video clips involving multiple consecutive episodes have started to find evidence that brain activity is naturally structured into events organized along these representational dimensions (e.g., Baldassano et al., 2017; Bird, 2020; Reagh and Ranganath, 2021; Lee and Chen, 2021; Heusser et al., 2021). Thus, if memory reactivation at episodic offset is an important neural signature of the rapid - ‘one-shot’ - learning of an unfolding realistic episode in humans, it is important to clarify whether it is concomitant to the encoding of episodes that included sequence of elements depicting coherent relations between them.

To address the issue, we designed a task that required participants to encode and later recall sequences of pictures depicting unique episodic-like events followed by a delay period with no stimulus. We used representational similarity analysis of scalp electroencephalography (EEG) recordings during encoding and found evidence for memory reactivation of the sequence elements of the episode after encoding, i.e., at the offset of the episode, and the degree of memory reinstatement at the offset predicted later memory recall for the specific episode. Memory reinstatement was not observed between successive elements within an episode, indicating that memory reactivation was specifically induced once participants perceived the unfolding episode to be completed. In a separate experiment, we also found that offset memory reinstatement was not present when participants encoded sequences of pictures that were not perceived as meaningful episodes. These results suggest that rapid memory reinstatement at episodic offset may be a neural signature engaged to integrate elements of the unfolding experience into a coherent memory for the just encoded episode.

## Materials and Methods

### Participants

Participants were native Spanish speakers who were recruited for pay (10€/h). All participants had normal or corrected-to-normal vision and reported no history of medical, neurological or psychiatric disorders. Twenty-five participants (17 females, age range 18-29) were recruited for Experiment 1. In addition, twenty-eight new participants (15 females, age range 19-35) were recruited for Experiment 2. Informed consent was obtained from participants in accordance with procedures approved by the Ethics Committee of the University of Barcelona.

### Experimental Procedure

Both Experiments (Fig. 1) consisted of an encoding and a retrieval phase, separated by a 10-15 mins-break in the middle. Task timing and visual stimulus presentation were under the control of commercially available software e-Prime 2.0 (Psychology Software Tools).

**Figure 1.**
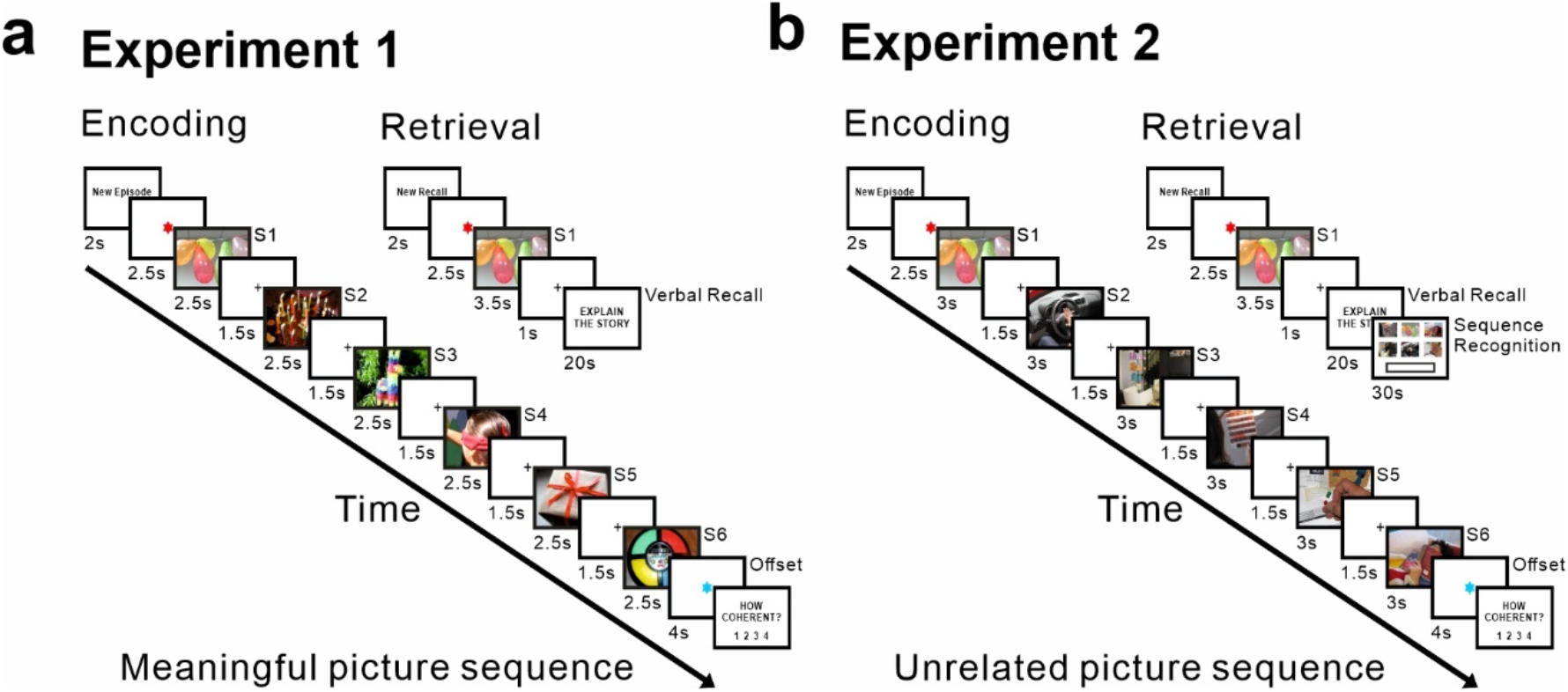
The experimental design. ***a***, Task design in Experiment 1. During encoding, 100 different sequences were presented only once. Each sequence included 6 different pictures that unfold a life-like coherent narrative episode. Each picture was presented 2.5s, followed by a 1.5s fixation cross. After each sequence of images, there was an offset period (4s) during which participants were instructed to avoid rehearsing the just-encoded picture sequence. Participants were asked to provide a subjective rating of episodic coherence to the just encoded sequence at the end of each trial. Retrieval task was conducted 10-15 minutes after encoding. During retrieval, the first picture of each sequence was presented for 3.5 s which was followed fixation cross, and a message prompted at the screen instructing to report the associated episode during encoding. Participants were asked to verbally report within 20 seconds their memory associated episode or to indicate whether no memory came up associated to that picture cue. ***b***, Task design in Experiment 2. Pictures were shuffled across sequences so that no meaningful story could be constructed after each sequence presentation. 60 shuffled sequence series were selected. The procedure was identical to Experiment 1 except two adjustments for task difficulty: 1. Time duration for the presentation of each picture during encoding was increased to 3000ms; 2. After each cued-picture recall task, participants were requested to perform a sequence order recognition task within 30 seconds, during which all 6 pictures from the same sequence series (including the cue picture) were presented on the screen in random positions and participants were asked to type the order of them as the original sequence presented during encoding phase.

The encoding phase of Experiment 1 included 100 different sequences, each consisting of 6 pictures that each was presented to participants only once. All pictures depicted emotionally neutral events and were controlled for saliency. All sequences of pictures described day life routine circumstances such as go shopping, reading a book at home or frying an egg in the kitchen. The pictures of each of the series were related to each other as in a story presented chronologically ordered. The order of pictures within each sequence was the same across participants but the order of the sequence presentation was counterbalanced across participants. Prior to the Encoding phase, participants were taught to attend to the picture series for a subsequent cue-recalled test yet not to engage in active rehearsal, especially when the blue fixation cross at the end of the series appeared. They were informed beforehand about the format of the subsequent test, entailing cued-recall of the images of each series. Participants were encouraged to memorize as many pictures as possible in a narrative form. The experiment only began when the examiner made sure that the participants fully understood the task. During the task, participants were encouraged to minimize eye movements and blinking. Each trial began with the presentation of text ‘New Episode’ for 2000ms, which marked the start of a new sequence. This was followed by a fixation screen with a red asterisk lasting 2500ms. Each picture was then presented sequentially on a white screen for 2500ms and followed by a 1500ms black fixation cross.

Immediately after the presentation of the last picture in each series, a blue asterisk was presented on the screen, indicating a post-episode offset period of 4000ms during which participants were previously instructed to avoid rehearsing the just-encoded picture sequence. The asterisk remained visible on the screen during the offset period. Immediately after the offset period, participants were asked to provide a subjective rating of coherence for the just encoded sequence. A rating scale ranged from 1 to 4, where 1 stood for ‘not coherent’ and 4 stood for ‘very coherent’. The next trial began after a fixed time interval of 2000ms.

For the retrieval phase, each trial started after the presentation of text ‘New Recall’ for 2000ms on the screen. This was followed by a fixation screen with a red asterisk lasting 2500ms. After the asterisk, the first picture of one sequence was presented to participant for 3500ms on the screen serving as a cue to prompt the free verbal recall for the rest of pictures in that sequence. Participants were instructed to start the verbal recall once the text ‘Explain the story’ was presented on the screen following a 1000ms fixation cross. The verbal recall was limited to 20 seconds and participants could stop the recall when finished by pressing the space bar. The order of the picture cues was randomized before their presentation at the retrieval phase.

In Experiment 2, we shuffled the pictures across sequences from Experiment 1 so that each sequence was formed by 6 pictures from different sequences. Thus, each sequence no longer depicted a meaningful episodic sequence. As in Experiment 1, the order of pictures within each sequence was kept the same across participants but the order of the sequence was randomized between participants. The general experimental settings for the encoding and retrieval phases and the instructions to participants were identical to Experiment 1. However, three adjustments were made: i) the presentation time of each picture during encoding was 3000ms; ii) the total number of sequences presented to the participants was sixty; and, iii) we added an order recognition task after each cued-picture recall task. During the sequence order recognition task, all 6 pictures from the same sequence (including the cue picture) were presented on the screen in random positions and participants were asked to type the order in which they appeared in the encoding phase. Participants had 30 seconds to type the order of the pictures and they could skip to the next trial when finished by pressing the space bar. These changes were motivated by previous pilot studies with small sample of participants that ensured this number of sequences provided a balanced outcome between picture sequences whose order memory was relatively preserved or not.

### EEG Recording

For both Experiments, EEG was recorded using a 31-channel system at a sampling rate of 500 Hz, using a BrainAmp amplifier and Ag/AgCl electrodes mounted in an electrocap (EasyCap) located at 28 standard positions (Fp1/2, Fz, F7/8, F3/4, Fc1/2, Fc5/6, Cz, C3/4, T3/4, Cp1/2, Cp5/6, Pz, P3/4, T7/9, P7/8, O1/2, Oz) and at the left and right mastoids. An electrode placed at the lateral outer canthus of the right eye served as an online reference. EEG was re-referenced offline to the linked mastoids. Vertical eye movements were monitored with an electrode at the infraorbital ridge of the right eye. Electrode impedances were kept below 3 kΩ. A band-pass filter (0.1 Hz -20 Hz) was implemented offline before the analysis.

### Verbal recall analysis

During the retrieval phase of Experiment 1 & 2, participants were asked to verbally recall in the form of a “narrative” as many pictures as possible within each sequence corresponding to the cue. Free verbal recall of each trial was recorded through an audio recorder and the audio files were later analyzed by a native Spanish speaker in the laboratory. Within each retrieval trial, a picture was considered as successfully retrieved when the precise details of the picture were described, or its core feature was mentioned during recall. Memory for each sequence was then quantified by the number of pictures (excluding the cue) correctly recalled.

### Sequence order analysis

For Experiment 1, the order of the verbally recalled items in each trial was analyzed. A trial was considered in-order when all recalled items followed the same sequential order as the encoding sequence. For Experiment 2, the temporal order memory for picture sequences recognition was compared to the true order of the sequence, and the result for each trial was coded as the maximum number of pictures (including the cue) correctly ordered in a row.

### EEG data analysis

For each participant, we first used EEG data of the encoding phase to extract epochs for each item within the sequence, namely an EEG epoch for the 1^st^, 2^nd^, 3^rd^, 4^th^, 5^th^ and 6^th^ picture. Each epoch had a duration of 2500ms and was baseline corrected to the pre-stimulus interval (−100 to 0ms). We then extracted epochs for the post-episode offset signals after each sequence with duration of 4000ms and baseline corrected to the time interval (−100 to 0ms) before its onset. Finally, we repeated the procedure to extract epochs for the post-item offset of 1500ms (with baseline corrected to -100 to 0ms), that corresponded to the inter-stimulus interval separating each item presentation during episodic sequence encoding phase.

### Time-resolved Representational Similarity Analysis (RSA)

For RSA, each EEG epoch data was smoothed by averaging data via a moving window of 100ms throughout the EEG epoch (excluding the baseline period) and then down-sampled by a factor of 5. RSA was performed at individual level and included spatial features (i.e., scalp voltages from all the 28 electrodes) (Silva et al., 2019). The similarity analysis was calculated using Pearson correlation coefficients, which are insensitive to the absolute amplitude and variance of the EEG response.

For both Experiment 1 & 2, we conducted a trial-based RSA between the EEG signal elicited by each encoding item (1^st^, 2^nd^, 3^rd^, 4^th^, 5^th^ and 6^th^) and the EEG signal elicited during immediate post-episode offset. After data smoothing and down-sampling, EEG epoch data for each item encoding contained 250 time points (given the 500 Hz EEG recording sampling rate) covering the 2500ms of item picture presentation, and EEG data for each post-episode offset contained 400 time points, equivalent to 4000ms. Point-to-point correlation values were then calculated, resulting in a single trial 2D similarity matrix with the size of 250×400, where the *x*-axis represented the offset time points and the *y*-axis represented the encoding time points. A trial-based RSA was computed between EEG patterns elicited by the encoding of 1^st^ to 4^th^ and 1^st^ to 5^th^ picture and the EEG patterns elicited at the immediate post-4^th^ and post-5^th^ picture ISI interval (i.e., 1500ms), respectively. More concretely, we conducted RSA between EEG signal elicited at post-stimulus period after the 4^th^ item and EEG patterns triggered during the encoding of each of the preceding 1^st^, 2^nd^, 3^rd^ and 4^th^ items. Then we repeated the same RSA but between EEG signal elicited at post-stimulus period after the 5^th^ item and EEG patterns triggered during the encoding of each of the preceding 1^st^, 2^nd^, 3^rd^, 4^th^ and 5^th^ items. This resulted in 9 similarity matrices in total for each participant. The resulting similarity matrices were then averaged which resulted in a single 2D matrix with size of 250×150 (i.e., 2500ms of picture encoding x 1500ms of post-stimulus offset or ISI), depicting the overall degree of similarity between EEG patterns elicited during item encoding and post-item offset.

### Nonparametric Cluster-Based Permutation Test

To account for RSA differences between conditions, we employed a nonparametric statistical method (Maris & Oostenveld, 2007), which identifies clusters of significant points on the resulting 2D similarity matrix and corrects for multiple comparison based on cluster-level randomization testing to control the family-wise error rate. Statistics were computed on values between conditions for each time point, and based on adjacency in the 2D matrix, adjacent points that passed the significance threshold (p < 0.05, two-tailed) were selected and grouped together as a cluster. The cluster-level statistics were then calculated by summing up the statistics of all time points within in each identified cluster. The procedure was then repeated 1000 times with randomly shuffled labels across conditions to simulate the null hypothesis. For each permutation, the cluster-level statistics with highest absolute value was registered to construct a distribution of the cluster-level statistics under the null hypothesis. The nonparametric statistical test was calculated by the proportion of permuted test statistics that exceeded the true observed cluster-level statistics.

### Bayes Factor statistical analysis

To further evaluate the power of the RSA effects that could be observed between conditions, we implemented the Bayes Factor statistical analysis (Kass & Raftery. 1995) in the point-to-point 2D similarity matrix. Bayes Factor was computed using Matlab Toolbox (Bart Krekelberg (2021). bayesFactor. https://github.com/klabhub/bayesFactor) on RSA values between conditions for each point of the resulting 2D similarity matrix (250×400). A Bayer Factor greater than 10 indicates strong evidence for difference between conditions comparing to the null hypothesis.

### Linear-mixed effect model

To investigate the relationship between EEG similarity values and behavioral memory on a trial basis we implemented a Linear Mixed Effect Model (LMM), which accounts for intra- and inter-individual variances. This analysis would also allow scrutinize the extent to which possible differences in EEG similarity results when comparing High and Low accuracy trials observed with a median-split approach described above were independent on the partitioning trial strategy implemented at subject level. Thus, we specified in our LMM the correlation values for one specific point on the resulting 2D similarity matrix as the dependent variable and included the following factors as fixed effect variables: the number of items correctly recalled (which ranged from 0 to 5, without counting the picture cue present at the recall task); the index of an item’
ss order in each sequence (1^st^, 2^nd^, 3^rd^, 4^th^, 5^th^ and 6^th^), and the coherence rating provided by the participant to each sequence at encoding (which ranged from 1 to 4). Participant number was introduced into the model as the grouping variable, with random intercept and a fixed slope for each of the fixed-effect variables. To balance the requirement for computational power and signal to noise ratio, we further smoothed the resulting 2D similarity matrix for each item-offset pair by averaging over a moving window of 200ms and then down-sampled by the factor of 5, both smoothing and down-sampling were conducted 2-dimensionally across the *x* and *y* axes. We applied the model fitting analysis independently for each position on the resulting 2D similarity matrix (50×80), then returned the 2D statistics map of the same size for each fixed-effect variable. Here to control for multi-comparison problem, the nonparametric cluster-based permutation test cannot be applied because each permutation represents a sample from the null distribution, which is not the case in LMM where it contains a covariance structure induced by multiple levels of relatedness among the individuals. Therefore, we implemented a Bonferroni correction to the statistical threshold to correct for the multiple comparison problem in the resulting statistical maps for each fixed-effect variable. Thus, the resulting statistical map was thresholded with an adjusted alpha level of α = 1.25×10-5 (0.05/4000).

## Results

### Experiment 1: Meaningful episodic sequence encoding

#### Free recall for meaningful episodes

In Experiment 1, participants were able to recall on average 2.32 items (SD = 0.456) in each series out of the total possible 5 items included in episodic sequence. Participants tended to recall encoded sequences in the form of “narrative” (e.g., “this is a party, there are balloons, the cake, and after the cake a piñata is broken and a gift comes out, but I don’t remember what it was”) and we counted the number of picture items included in their recall. The mean percentage of trials across participants that successfully recalled 0,1,2,3,4 and 5 items after the retrieval cue were respectively 24.89% (SD = 9.34%), 8.65% (SD = 3.41%), 14.14% (SD = 4.94%), 24% (SD = 5.68%), 18.91% (SD = 6.49%) and 9.41% (SD = 6.35%). A repeated-measures ANOVA revealed that participants’ memory recall differed as a function of number of items recalled following the retrieval cue (*F*(5,120) = 26.227, p < 0.001)

(Fig. 2*a*). To increase the signal to noise ratio, for later RSA on EEG data, we first adopted a median split approach to separate the trials based on the corresponding task performance. Sequences with 2 or fewer items recalled during the retrieval phase were labeled as Low memory trials and sequences with equal or more than 3 items recalled were labeled as High memory trials. The threshold was selected by its relatively well-balanced separation for number of trials at subject level, as the average percentage of trials after the median split separation was respectively 47.64% (SD = 12.51%) for Low memory condition and 52.36% (SD = 12.51%) for High memory condition (Wilcoxon signed-rank test: *z* = 0.821, *p* = 0.412) (Fig. 2*b*).

**Figure 2.**
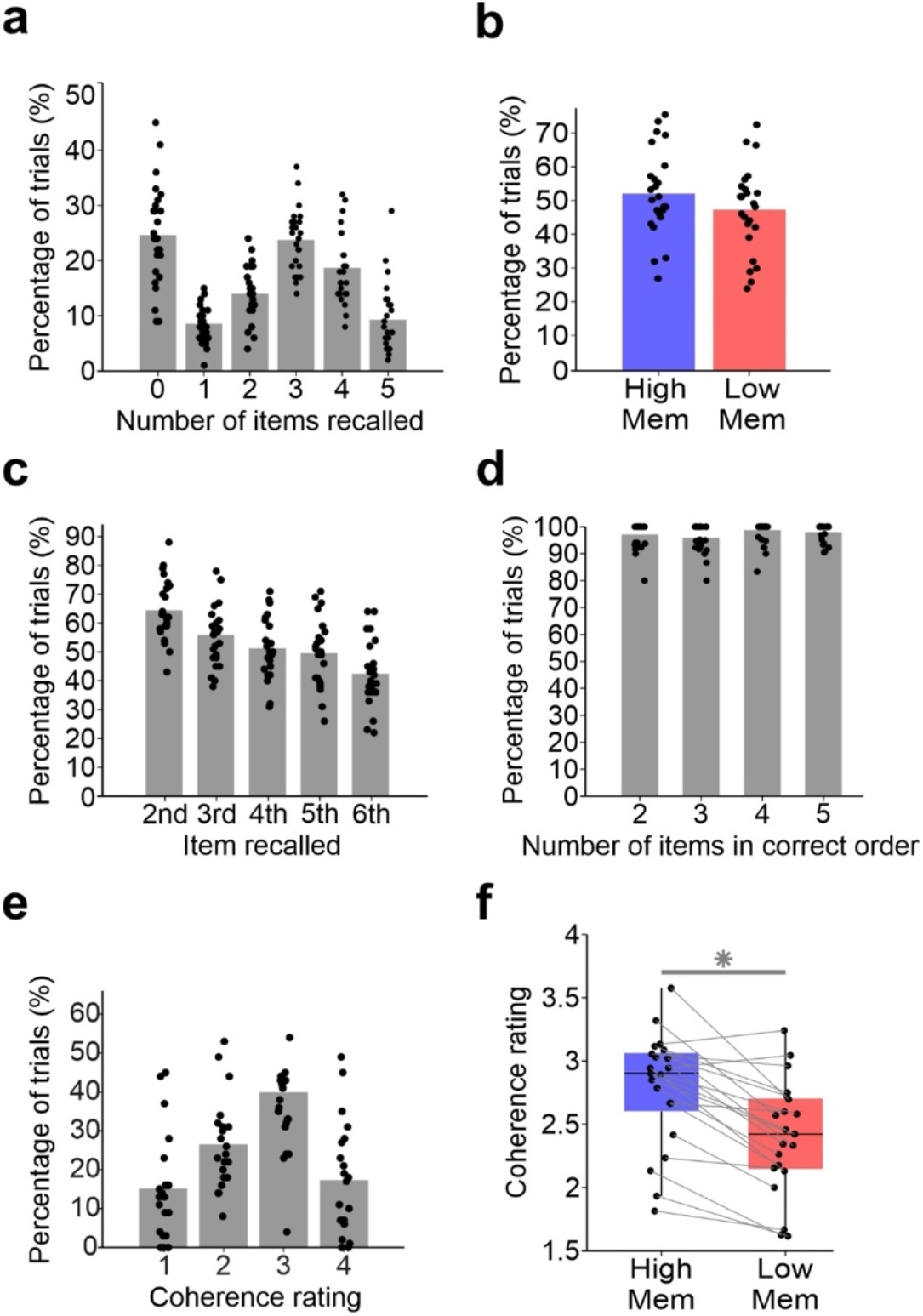
Behavioral results for Experiment 1. ***a***, Percentage of trials at the cued recall task as a function of number of items recalled in each sequence. ***b***, Percentage of trials included in the High (High Mem) and Low memory (Low Mem) condition in experiment 1 after implementing the median-split approach. Series with at least 3 item pictures correctly retrieved after cue were counted as High memory trials, otherwise the series were counted as Low memory trials. ***c***, Percentage of trials as a function of the item across sequence order being recalled. ***d***, Percentage of trials with recall in-order as a function of number of items recalled. A recall trial was considered as in-order when its items were recalled following the same sequential order as the true sequence. ***e***, Percentage of trials as a function of the participants’ degree of subjective coherence rating. ***f***, Mean coherence rating score for trials included in the High and Low memory condition. separated by median-split verbal recall memory. In (a-c), bars represent the average across participants. Each black dot represents values for an individual participant. For all boxplots in (d), the central mark is the median, the edges of the box are the 25th and 75th percentiles. \**p*< 0.05.

For each participant, we then calculated recall accuracy for each of the items in the sequence. The results showed a gradual decrease in accuracy for items as a function of their order position in the sequence during encoding: mean = 65.09% and SD = 10.44% for item 2, mean = 56.48% and SD = 10.92% for item 3, mean = 51.76% and SD = 10.43% for item 4, mean = 50.06% and SD = 11.57% for item 5 and mean = 42.91% and SD = 11.35% for item 6 (repeated-measures ANOVA: *F*(4,96) = 96.475, p < 0.001) (Fig. 2*c*). We also assessed how well item order was preserved during recall by counting, for each trial sequence, the number of items recalled in correct order as a function of the total number of recalled items for that sequence. We found that participants were accurate in recalling in order the items, independent of the total number of items included in their recall (*F*(3,71) = 1.464, p = 0.231; mean = 97.17% and SD = 5.40% for 2 items, mean = 95.89% and SD = 5.51% for 3 items, mean = 98.79% and SD = 5.36% for 4 items, mean = 98.01% and SD = 3.26% for 5 items) (Fig. 2*d*).

#### Subjective ratings of episodic coherence

The coherence rating provided a subjective measure of the degree of perceived narrative of each sequence. Due to technical issues, data for coherence ratings of 4 participants were not registered and they could not be included in the analysis. On average, for the remaining 21 participants, sequences were rated as 2.6 (SD = 0.41) (on a scale that ranged from 1 to 4), and the mean percentage of trials rated as 1, 2, 3, and 4 were respectively 15.34% (SD = 13.63%), 26.79% (SD = 11.67%), 40.36% (SD = 17.04%) and 17.51% (SD = 14.58%) (*F*(3,60) = 9.889, *p* < 0.001) (Fig. 2*e*). After median splitting the trials based on verbal recall performance, trials with High memory showed significantly higher coherence ratings (mean = 2.79, SD = 0.45) compared to trials with Low memory (mean = 2.40, SD = 0.45; paired Student *t*-test: *t*(20) = 6.166, *p* < 0.001, two-tailed) (Fig. 2*f*).

#### RSA between item sequence and episodic offset at encoding

We first asked whether EEG patterns induced at the post-episode offset period correlated to EEG patterns elicited by the just encoded picture items within the episodic sequence, and if so, whether the magnitude of such correlation was associated to memory recall at the test. To address this issue, we implemented a trial-based RSA between EEG data elicited at picture item encoding (1^st^, 2^nd^, 3^rd^, 4^th^, 5^th^ and 6^th^) with EEG data at the immediately following episode offset period. For each participant, the resulting trial-based RSA values were averaged separately according to two memory conditions: those associated with the participants ability to recall picture items with High memory (3 or more items) and those with Low memory (less than 3 items). To account for enough number of EEG trials to be included in both conditions and to ensure these trials included all items (1-6) and post-episode offset EEG signal cleaned of artifacts, we set a post-hoc criteria to exclude participants that did not reach a minimum of >15% number of trials in either condition, resulting in 15 participants for the current analysis (Average percentage of trials remained well-balanced between conditions with 35.04% (SD = 8.06%) of total trials included in analysis for Low memory condition and 32.75% (SD = 11.29%) of total trials for High memory condition; Wilcoxon signed-rank test: *z* = 0.597, *p* = 0.551).

In both High and Low memory conditions, the results of this analysis revealed an increase in similarity between EEG patterns induced ∼400ms - 800ms at the post-episode offset period and EEG signal elicited ∼400ms - 1300ms at picture item sequence encoding period (Fig. 3*a*). However, the nonparametric cluster-based permutation analysis identified one statistically significant cluster where similarity values were higher in the High than in the Low memory trials (*p* = 0.001; mean t-value = 3.242, peak t-value = 5.473) (Fig. 3*b*). We next evaluated whether the similarity between item encoding and post-episode offset period was driven by EEG patterns elicited by specific picture items within the just encoded sequence. We extracted the mean similarity values within the identified cluster for each item-offset pair and computed a repeated-measures ANOVA with two factors: trial condition (High vs Low memory) and encoding item (1^st^, 2^nd^, 3^rd^, 4^th^, 5^th^ and 6^th^). The results of this analysis showed a significant main effects for both trial condition (*F*[1,14] = 15.407, *p* = 0.002) and encoding item (*F*[5,70] = 2.677, *p* = 0.028), but no significant interaction (*F*[5,70] = 0.315, *p* = 0.902). (Fig. 3*c*), indicating that episodic offset increase in similarity was not driven by EEG patterns elicited by specific items from the encoded item sequence. To further evaluate the power of the effect of the difference between High and Low memory conditions, we calculated the Bayes Factor on similarity values of each point on 2D similarity matrix. The results showed strong evidence (Bayes Factor greater than 10) for difference between conditions overlapping the area where significant higher similarity values were found on nonparametric cluster-based permutation analysis (Fig. 3*d*).

**Figure 3.**
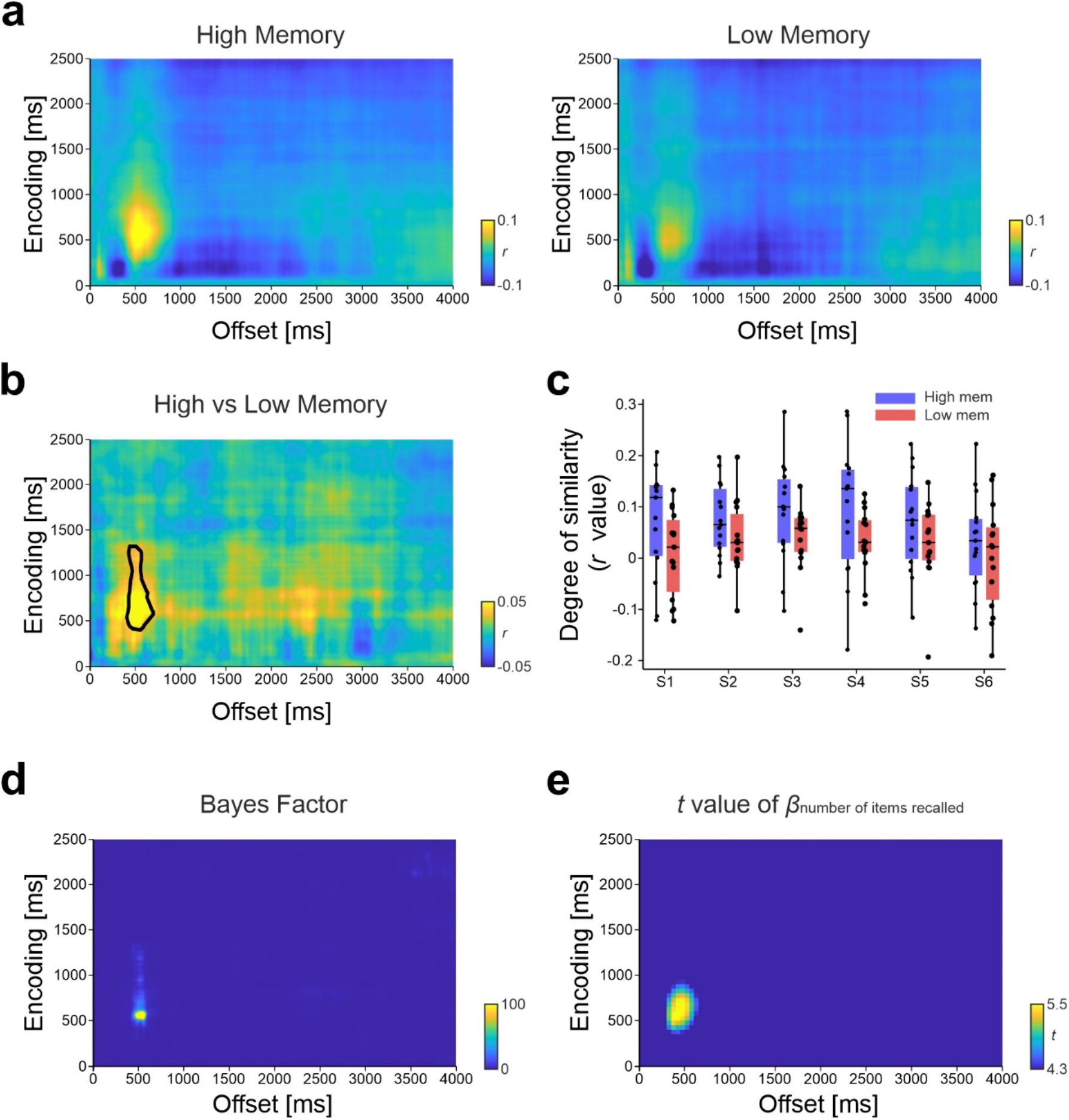
Neural similarity results at post-encoding period for Experiment 1. ***a***, Time-resolved degree of neural similarity between item picture encoding and post-encoding offset for events with High or Low memory at test. ***b***, Difference between similarity values for the two conditions. Statistically significant (*p* < 0.05, cluster-based permutation test) higher similarity value ∼400ms - 800ms at the post-episode offset period with ∼400ms - 1300ms at item picture encoding period was found for events with higher recall performance (indicated by a black thick line). ***c***, In all boxplots the central mark is the median across participants, the edges of the box are the 25th and 75th percentiles. They depict the degree of similarity within the identified cluster for each item encoding with its corresponding offset period. Each black dot represents values for an individual participant. High memory sequence showed significantly greater similarity across encoding items. ***d***, Bayes Factor of the difference between similarity values for the two conditions. A Bayes Factor greater than 10 indicates strong evidence for difference between High and Low memory condition. ***e***, *t*-value map of the variable numbers of item recalled reveals the area that exceeded the significance threshold after Bonferroni correction with adjusted alpha level of α = 1.25×10^−5^ (one-tailed). No area exceeding the significance threshold was found for *t*-value map of the variable serial position of item in sequence neither for that of the variable coherence rating.

Having shown that neural similarity increases of the just encoded sequence elements was elicited at episodic offset and that it was functionally associated to later memory recall, we then leveraged this to explore the relationship between the magnitude of memory reinstatement and the numbers of items to be recalled correctly later. At the same time, we asked whether the observed effects could be simply explained by the participants’
s subjective feeling of coherence of the episode, as we found that subjective ratings of coherence were higher for High than for Low memory condition. To address these issues, we applied a LMM to our trial-based RSA data (see Methods). Two participants from the previous RSA analysis were not included here due to the missing data for coherence rating, resulting in 13 participants in total for the LMM analysis. Given that previous median-split analysis showed an increase similarity magnitude associated with higher recall performance, we specifically focused on this trend for LMM analysis. The result revealed one time interval that survived the statistical threshold for the fixed effect variable total number of items correctly recalled (one-tailed, mean *t*-value = 5.032, peak *t*-value = 5.621, *p* < 1.25×10^−5^, Bonferroni corrected) (Fig. 3*e*). The time interval, which covered ∼300ms - 700ms of post-episode offset and ∼300ms - 900ms of item encoding, indicated the region where the degree of neural similarity of each item elicited during post-episode offset was significantly positively correlated with total number of items to be recalled in the corresponding sequence. However, no significant point exceeding the statistical threshold was identified on the statistical map accounting for the variable indexing order position in sequence nor for the variable indexing coherence rating.

#### RSA between item encoding and sequence item immediate offset

Next, we asked whether the increases in neural similarity between EEG patterns elicited at item sequence encoding and encoding offset were specific to post-episode period, or alternatively, whether they could be also found at offset periods immediately following picture encoding. To address this issue, we implemented the RSA between EEG pattern elicited by each item in the encoding sequence and the EEG signal pattern induced during the immediate post-item offset period. The current analysis was centered in the post-item offset period after the 4^th^ and 5^th^ item as this represented a delay period, as in episodic offset period, that is preceded by the encoding of multiple items from a sequence but differed in that the encoding of episode is not completed yet. At the same time, this research strategy allowed us to implement the same median split analysis used in our previous analysis (i.e., whether or not at least 3 items after the picture cue were correctly remembered), thereby enabling the comparison of the RSA results from the two conditions later on. 8 out of the total 25 participants were excluded for this analysis due to insufficient number of clean EEG trials in either condition (i.e., at least 15% of total number of trials in either encoding item of either condition) and the remaining participants maintained the balanced separation of trials between conditions with 41.06% (SD = 12.89%) of total trials included in analysis for Low memory condition and 37.44% (SD = 9.05%) of total trials for High memory condition (Wilcoxon signed-rank test: *z* = 0.355, *p* = 0.722). The result of this analysis showed no clear increases in neural similarity in either High or Low memory conditions (Fig. 4*a*) and that no cluster of similarity values were accounted when the two conditions were compared with a cluster-based permutation test (Fig. 4*b*). With the aim to further examine that the increase in neural similarity was specific to the episodic offset period, we directly compared the neural similarity findings at the first 1500ms of the episodic offset vs at the 1500ms of the 4^th^ and 5^th^ post-item offset. The results of this analysis confirmed a cluster of significantly higher neural similarity at the episodic offset condition (Fig. 4*c*), thereby corroborating the notion that the increase in neural similarity was specific to post-episodic encoding delay period.

**Figure 4.**
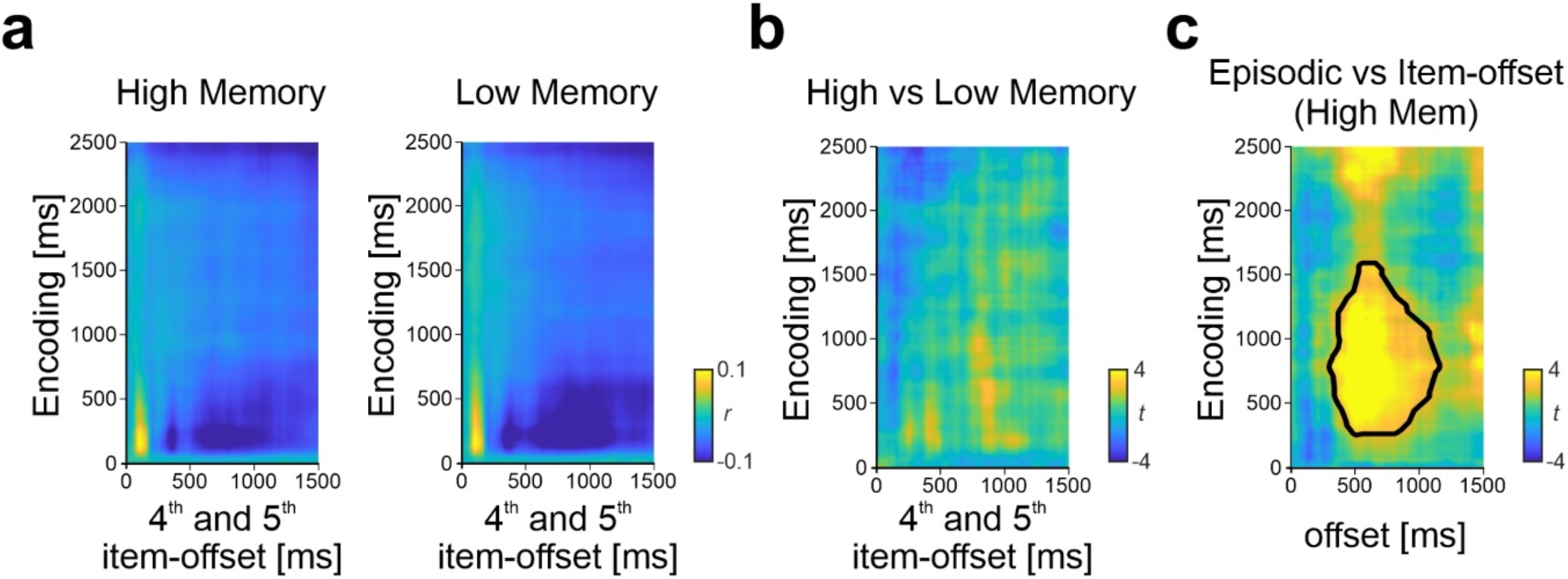
***a***, Time-resolved degree of similarity between item picture encoding and 4^th^ and 5^th^ post-item following the events that were later recalled with High or Low memory. ***b***, Difference (expressed in *t* values, uncorrected) between similarity values for the 4^th^ and 5^th^ post-item High and Low memory conditions. No cluster indicated significantly different similarity values between conditions (two-tailed, *p* < 0.05, cluster-based permutation test). ***c***, Difference (expressed in *t* values, uncorrected) between similarity values for episodic offset and the 4^th^ and 5^th^ post-item offset in the High memory condition. Higher neural similarity was found during the episodic offset compared to item-offset. The significant cluster is indicated by a black thick line. *p* < 0.05, corrected with a cluster-based permutation test.

### Experiment 2: Non-meaningful episodic sequence encoding

#### Behavioral results

In general, participants were able to recall on average 0.14 items (SD = 0.176) out of the possible five (picture cue was not included in the counting) in each series. The mean percentage of trials across participants to successfully recall 0,1,2,3,4 and 5 items after the cue were respectively 89.23% (SD = 11.92%), 7.98% (SD = 8.27%), 1.96% (SD = 3.14%), 0.77% (SD = 1.84%), 0.06% (SD = 0.32%) and 0% (SD = 0%) (*F*[5,135] = 795.913, *p* < 0.001) (Fig. 5*a*). Even though participants were unable to verbally recall almost any item from an encoded sequence, they showed above chance performance in the order item sequence recognition task.

**Figure 5.**
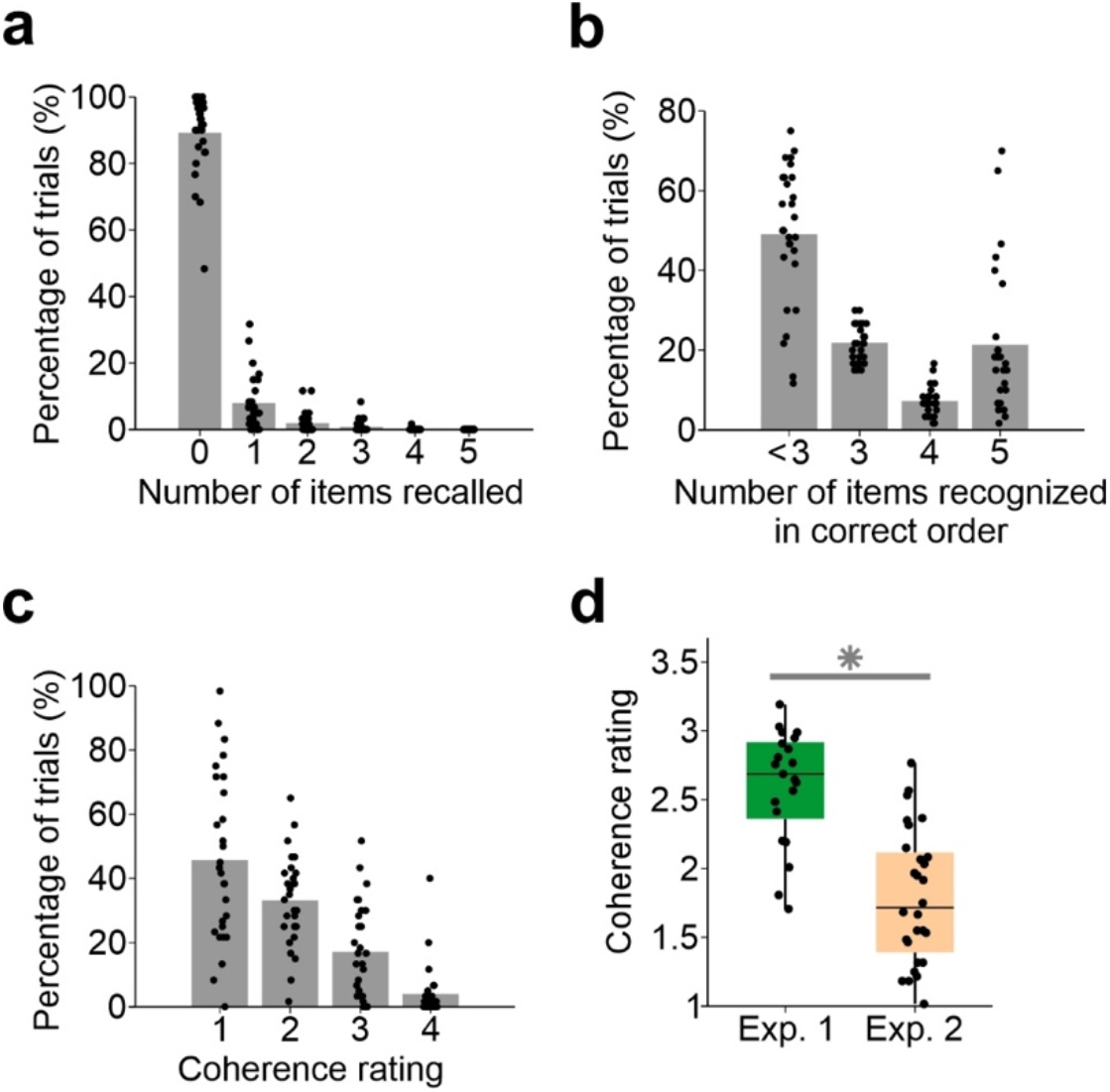
Behavioral results for Experiment 2 ***a***, Percentage of trials separated by number of pictures correctly recalled after the picture. ***b***, A temporal order sequence recognition memory test was added after the cued recall task. In the recognition memory task, all 6 pictures from the sequence were presented in random positions at the screen, including the cue, and participants had 30 s to order the correct temporal structure of the encoded sequences. The figure showed percentage of trials separated by the sequence recognition score. The score is quantified as the maximum number of pictures correctly ordered consecutively in one trial. A trial with less than 2 correct pictures consecutively ordered following either the cue, the 2^nd^, the 3^rd^ or the 4^th^ picture (score less than 3) was counted as no sequence recognized. ***c***, Percentage of trials separated by degree of coherence rating. ***d***, Mean coherence rating score of sequences in Experiment 1 vs Experiment 2. In (a-c), bars represent the average across participants. Each black dot represents values for an individual participant. For all boxplots in (d), the central mark is the median, the edges of the box are the 25th and 75th percentiles. \**p*< 0.05.

We analyzed these data by quantifying the maximum number of pictures correctly ordered consecutively for each given trial sequence. A trial was considered to have been recognized if the participant reported at least 3 items in correct order from the sequence. On average, the percentage of trials to have scored less than 3, equal to 3, 4 and 5 (i.e., all pictures in order following the cue) were respectively 49.11% (SD = 17.31%), 21.85% (SD = 4.72%), 7.26% (SD = 3.55%) and 21.79% (SD = 17.62%) (*F*(3,81) = 39.777, *p* < 0.001) (Fig. 5*b*). The participants provided correct order recognition for 50.89% (SD = 17.31%) of the trials, which is statistically significantly above chance (chance level = 15%, *t*(27) = 10.975; *p* < 0.001, two-tailed). Finally, the average coherence rating for sequences in Experiment 2 was 1.79 (SD = 0.48), and the mean percentage of trials rated as 1, 2, 3, and 4 were respectively 45.71% (SD = 25.96%), 33.15% (SD = 14.20%), 17.14% (SD = 14.68%) and 3.99% (SD = 8.31%) (*F*(3,81) = 24.108, *p* < 0.001) (Fig. 5*c*). In general, sequences in Experiment 1 were rated significantly higher than sequences in Experiment 2 (*t*(47) = 6.164, *p* <0.001, two-tailed) (Fig. 5*d*), which suggested that the subjective feeling of coherence matched the general manipulation of the experiment.

#### RSA between item sequence and episodic offset at encoding

As in experiment 1, we computed a trial-based RSA between 2500ms EEG patterns elicited by each sequence picture item and the corresponding EEG data induced at the 4000ms episodic offset. The parameters for data smoothing and down sampling was kept the same as Experiment 1. For each participant, the resulting 2D similarity matrix was first averaged within each item-offset pair and then across pairs. 10 out of the total 28 participants were excluded for this analysis due to insufficient number of clean trials for all items (at least 15% of total number of trials in each item-offset pair). Thus, a total of 18 participants were included in the analysis. However, and contrary to as in experiment 1, the result of RSA for Experiment 2 did not show any observable neural similarity increase at the offset period (Fig. 6*a*). To assess the extent to which neural similarity patterns seen in experiment 1 differed from those obtained in experiment 2, we separately compared the neural similarity values for High and for Low memory conditions in Experiment 1 with those obtained in Experiment 2. The results of these analyses revealed that neural similarity increase found at early offset period in experiment 1, for both High and Low memory conditions, was statistically different from similarity values at the offset period in experiment 2 (Fig. 6*b*). More concretely, this analysis returned one significant cluster (*p* = 0.022 (corrected), mean *t*-value = 3.354, peak *t*-value = 5.059), that comprised higher neural similarity values for EEG data within ∼400 - 1000ms time range from post-episode offset period and ∼200-1400ms time range from item picture encoding from High memory trials in experiment 1 over trials from experiment 2. A similar cluster in its timing (∼450 – 1000ms of post-episode offset and ∼200-1100ms of item encoding) with significantly higher neural similarity values was found when comparing Low memory trials from experiment 1 with trials from experiment 2 (*p* = 0.026 (corrected), mean *t*-value = 2.928, peak *t*-value = 4.237).

**Figure 6.**
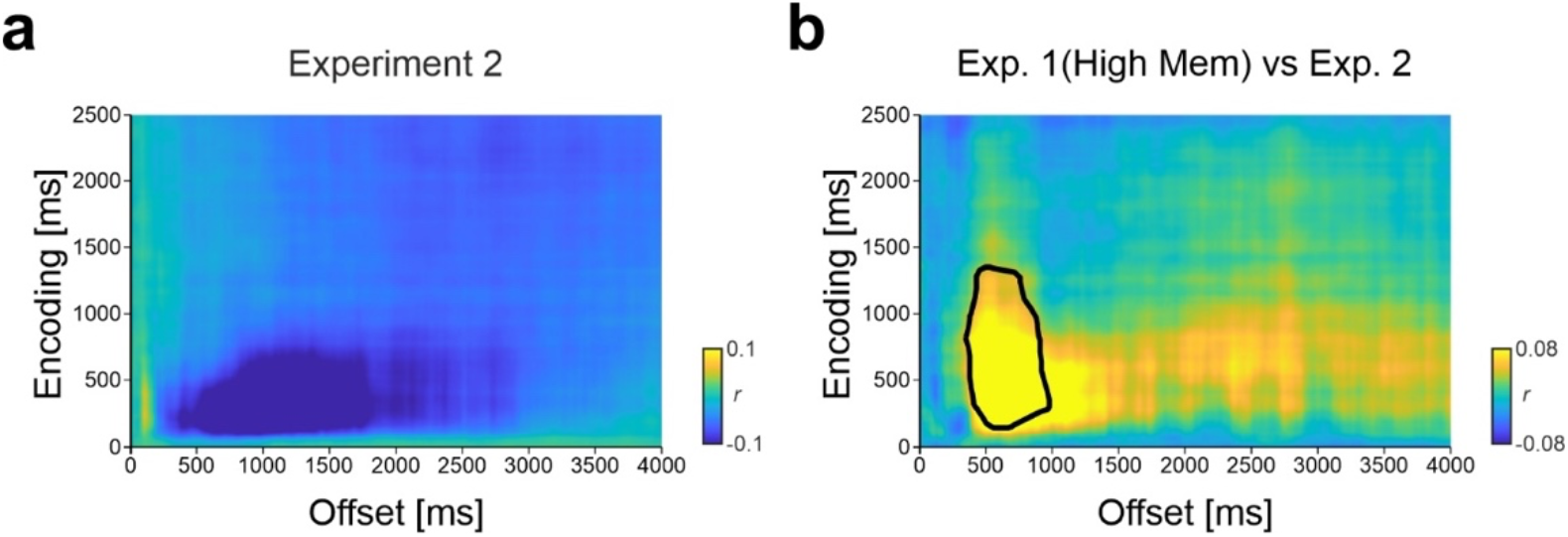
Neural similarity in Experiment 2. ***a***, Neural similarity between item picture encoding at post-encoding period for Experiment 2. ***b***, Higher neural similarity was found during the offset period in High memory trials in Experiment 1 when compared to the offset period in in Experiment 2 (the significant cluster is indicated by a black thick line. *p* < 0.05, corrected with a cluster-based permutation test).

Finally, we assessed whether the participants’ ability to preserve a temporal order memory, irrespective of their lack of episodic recall could still be associated to increases in neural similarity at the picture sequence offset period. For each individual, we split the encoding trials as a function of whether they were accurate in the temporal order recognition test (defined by trials with 3 items in correct order) and those trials where the participant showed poor temporal order accuracy (i.e., < 3 elements in correct order). In this analysis, 4 more participants were excluded due to insufficient number of clean EEG trials in either condition. The results of this analysis revealed no significant differences at cluster level (p > 0.05 two-tailed; permutation test) between the two conditions.

## Discussion

Here, we investigated in healthy human participants whether memory reinstatement of a just-encoded sequence of episodic items is a mechanism selectively engaged to support episodic memory formation. In two separate experiments, we examined whether memory reinstatement at encoding offset was concomitant to meaningful and/or to non-meaningful sequence of episodic items. In a first experiment, we used representational similarity analysis of scalp EEG recordings during the encoding of sequences of pictures depicting unique episodic-like events and we found that EEG patterns elicited during picture viewing correlated with EEG patterns at the episode offset. The degree of episodic offset-reactivation predicted later memory recall of the encoded picture sequence. In a second experiment, we used a similar analytical approach on EEG recordings while a different set of participants encoded sequences of pictures that were unrelated to each other, thereby preserving similar temporal encoding structure on meaningless episodic sequences. In this experiment, we did not find evidence of post-encoding memory reactivation at the offset. These results suggest that post-encoding memory reinstatement is akin to the rapid formation of unique and meaningful episodes that unfold over time.

Given the unfolding nature of our experience, researchers have started to focus the attention to brain encoding mechanisms that followed online encoding, as this may offer an “optimal” window whereby an unfolding episode can be registered as a bound representation once the ongoing inputs have concluded (Lu et al., 2020). fMRI studies using short videoclips (Zacks et al., 2001; Ben-Yakov et al., 2011; Ben-Yakov et al., 2013), sequence learning tasks alternating picture categories (DuBrow and Davachi, 2014) and, more recently, using long movie clips (Baldassano et al., 2017; Ben-Yakov et al., 2018) offered converging evidence that the brain is sensitive to episodic boundaries during encoding. In line with these findings, we recently showed that event boundaries, operationalized as transition points in the encoding time whereby one episode ends and new one starts, triggered the rapid memory reinstatement of the just encoded event information upon context shifts, and that the degree of memory reinstatement predicted the participants’ ability to preserve temporally adjacent events in a later test (Sols et al., 2017; Silva et al., 2019). The current results extend previous findings in several important ways.

If memory reactivation at episodic offset serves to promote the encoding of unique events into memory, we reasoned, then, it should be observable at the end of an episode and not contemporarily to the beginning of another episodic input. Our results from experiment 1 showed that this is the case, as EEG patterns elicited during sequence encoding correlated to EEG patterns at immediate offset periods that did not contain any stimuli input. Importantly, memory reinstatement was not observed in transition points between pictures of an episode. These findings lend support to the notion that memory reinstatement does not merely reflect an ongoing mechanism that links items associated during encoding. Instead, it suggests that episodic offset memory reactivation is a specific neural signature induced once an individual perceives an unfolding episode concluded.

We also found that post-encoding memory reinstatement followed the encoding of picture sequences that were perceived by individuals as depicting a meaningful episode (i.e., Experiment 1) but not after the encoding of sequences of pictures that were unrelated to each other (i.e., Experiment 2). Why would post-encoding reactivation be important for the formation of memories of meaningful episodes? Previous research has shown that the recall of life-like episodes is organized along representational dimensions beyond their temporal structure, such as the causal (Brownstein & Read, 2007), semantic (van Kesteren et al., 2013; Baldassano et al., 2018) and the relations between the elements embedded in the episodes (Lee and Chen, 2021). This research aligns well to psychological research that emphasized that memory is carved by how people construct a high-order model of the ongoing experience and that the detection of an episodic boundary triggers a set of neural and cognitive processes that would allow the integration of the just-encoded episodic model in memory (Radvansky and Zacks, 2011). Importantly, for this resulting model to be effectively recalled later, it should include several components that vary in the representational hierarchy: from object features to semantics (Radvansky and Zacks, 2011). Recent fMRI research has provided evidence that such representational structure is well reflected along cortical hierarchy during online encoding of realistic stream of stimuli (e.g., Baldassano et al., 2017; Bird, 2020; Reagh and Ranganath, 2021; Lee and Chen, 2021; Heusser et al., 2021) and that such cortical patterns are coupled to hippocampal activity at the detection of high-order event boundaries during encoding to account for later recall (Baldassano et al., 2017). Our current findings contribute to this literature by indicating that memory reactivation is a neural signature by which this high order episodic models can be stored rapidly at episodic encoding offset.

The fact that memory reactivation was found during a delay period immediately following encoding can be seen as a reflection of a mechanism inherently linked to the working memory (WM) maintenance of the encoded sequence of pictures. However, several observations in our results suggest the reinstatement at the episode offset cannot be explained solely by WM processes. First, post-encoding memory reinstatement predicted participants’ ability to recollect the episodic picture content in a later test but not their subjective rating of coherence that immediately followed the delay period. Should post-encoding memory reinstatement be a mechanism supporting WM maintenance we would expect it to be at least partially associated with the individual’s ability to evaluate the episodic coherence of the encoded picture sequence after the delay period. Second, post-encoding memory reinstatement in our study was circumscribed to the beginning but not throughout the post-encoding delay period. Studies investigating the neural substrates of WM have shown that delay maintenance is associated with a sustained increase in activation of neocortical structures (Fuster and Alexander, 1971). Should post-encoding memory reinstatement be associated with the sustained increase neural activity during the delay we would expect it to be observed over extended portions of the offset delay period and not only at the beginning of it. More recently, it has been argued that such above-threshold delay-period activity may support functions other than information storage per se (D’Esposito and Postle, 2015) and the existence of other neural coding mechanisms such as “activity-silent” states (Stokes, 2015) and dynamic coding schemes (Liu et al., 2020). However, these neural representational formats are still susceptible to be identified with the implementation of multivariate decoding approaches, such as the one implemented in the current design, thereby rendering unlikely they were unobservable throughout the delay period in our study. Third, recent findings from direct recordings at hippocampal and neocortical regions in epileptic human patients showed that the hippocampus marks the conversion from external (perceptual) to internal (mnemonic) representations by signalling cortical reinstatement at ∼500 ms after the onset of a retrieval cue (Treder et al., 2021). Our study showing that memory reinstatement detected from scalp EEG signal emerged transient and of a relatively brief duration at ∼ 500 ms at post-episodic encoding period may result from a switch from perception to memory process during encoding itself which would help bind the unfolding information into a memory episodic unit.

A pressing question derived from the findings of our second experiment is why post-encoding memory reactivation was absent when participants encoded sequences of pictures depicting unrelated content. Though the statistical absence of an effect cannot guarantee the absence of the effect, our results lend support to the notion that that post-encoding reactivation strength may not be an automatic mechanism that links items associated during encoding. Instead, it suggests that it contributes to the integration of event components into long-term memory once an individual perceives a meaningful episode is concluded. Previous fMRI research has highlighted that the primary role of hippocampal offset signal in reflecting binding operations of stimuli that just co-occurred within the same spatial-temporal context (Staresina and Davachi, 2009; Ritchey and Cooper, 2020). Similarly, fMRI studies (Baldassano et al., 2017) and electroencephalographic recordings from implanted electrodes in epileptic patients (Michelmann et al., 2021) revealed that the degree to which offset hippocampal activity couples with cortical patterns of activity during a continuous stream of stimuli predicts pattern reinstatement during later recall, thereby indicating that the hippocampus may be responsible for binding cortical representations into a memory trace online during encoding (McClelland et al., 1995; Norman and O’Reilly, 2003; Moscovitch et al., 2005).

Single-cell recordings from the rodent hippocampus during navigational tasks have shown that neural replay can be observed after the first lap on a novel track (Foster and Wilson, 2006). More strikingly, this research has shown that post-encoding replay may preserve the temporal structure of the encoded event sequence in a compressed time-manner (Csicsvari et al., 2007; Diba & Buzsaki 2007; Foster & Wilson 2006; Gupta et al., 2010; Karlsson and Frank, 2009), thereby suggesting that awake neural replay after single-shot learning may reflect the encoding of a model of the experience in long-term memory (Foster et al., 2017). Our findings based on scalp EEG recordings are blind to whether memory reinstatement at episodic offset period relies on memory replay of a temporally preserved structure of the encoded sequence. Future studies using brain acquisition approaches more sensitive to hippocampal activity, such as MEG (e.g., Liu et al., 2019) or intracortical recordings directly from the human hippocampus (e.g., Vaz et al., 2020) may help disambiguate whether the compressed episodic offset memory reinstatement preserves a temporal structure of an encoded sequence episode.

To conclude, we have shown that episodic offset memory reinstatement is selectively engaged to support successful encoding of sequential picture series with a coherent structure. These results shed light on the neural mechanisms that support the rapid learning of novel episodes that unfold over time in humans and how they serve to selectively transform experiences into long-term memory representations that can be later recalled.

## Acknowledgments

We thank Íria Rodríguez for assistance in data collection and David Cucurell in technical assistance. X.W. is supported by a PhD fellowship from the University of Barcelona. B.P.S. is supported by a Wellcome Trust/Royal Society Sir Henry Dale Fellowship (107672/Z/15/Z). A.B.Y is supported by a Dorothy Hodgkin Royal Society Fellowship (DHF\R1\191141). L.F. is supported by Ministerio de Ciencia e Innovación, which is part of Agencia Estatal de Investigación (AEI), through the project PID2019-111199GB-I00 (Co-funded by European Regional Development Fund. ERDF, a way to build Europe) and by ICREA Academia. We thank CERCA Programme/Generalitat de Catalunya for institutional support.

